# Roosting ecology of the southernmost bats, *Myotis chiloensis* and *Histiotus magellanicus*, in southern Tierra del Fuego, Chile

**DOI:** 10.1101/2020.04.29.068130

**Authors:** Gonzalo Ossa, Thomas M. Lilley, Austin G. Waag, Melissa B. Meierhofer, Joseph S. Johnson

**Affiliations:** ConserBat EIRL, San Fabian, Chile; Programa para Conservación de los Murciélagos de Chile, Santiago, Chile; Finnish Museum of Natural History, University of Helsinki, Helsinki, Finland; Department of Biological Sciences, Ohio University, Athens, Ohio, USA; Department of Rangeland, Wildlife and Fisheries Management, Texas A&M University, College Station, Texas, USA

**Keywords:** Chilean myotis, day-roosts, Karukinka, southern big-eared brown bat, thermoregulation, torpor

## Abstract

There are few studies of day-roosting ecology of bats inhabiting the southernmost forests of South America, where cool summer temperatures and land management practices pose several challenges. The goal of the present study was to describe day-roosting habitats and patterns of thermoregulation in two bat species occurring on Tierra del Fuego, *Myotis chiloensis* (Chilean myotis) and *Histiotus magellanicus* (southern big-eared brown bat), during late spring. To do so, we tagged 17 bats with temperature-sensitive radio-transmitters, located 17 day-roosts, and collected 81 days of skin temperature data. We concurrently recorded ambient air temperature to determine its effect on torpor use. Both species were found roosting in large diameter (77.8 ± 6 cm), typically live, *Nothofagus pumilio* trees (lenga) located on the edges of forest gaps or within stands primarily composed of smaller, younger trees. Bats of both species frequently used torpor, with skin temperatures dropping below a torpor threshold on 89% of days (*n* = 72) and daily minimum skin temperatures averaging 16.5 °C over the course of our study. Average daily air temperature was a significant predictor of torpor use, with lower skin temperatures and more time spent in torpor observed on colder days. Minimum skin temperature and time spent torpid did not vary between bat species, nor did the characteristics of day-roosts. These data show that spring temperatures in Tierra del Fuego pose an energetic challenge that bats meet through frequent use of torpor, and likely, habitat selection. We recommend local conservation efforts keep these thermal challenges in mind by retaining large trees, which may provide warmer microclimates or room for social groups.

## INTRODUCTION

Many bat species rely solely on forests for roosting and foraging habitats. The relationships between bats and these habitats are complex, with species using forested areas non-randomly based upon their unique morphologies, behaviors, and physiologies (Fenton & Bogdanowicz 2002; Fabianek *et al*. 2015, Vasko *et al*. 2020) and the structure and composition of the vegetative community (Hayes 2007; Law *et al*. 2016). For example, several species roosting in cavities or under bark select the tallest and largest diameter trees available, and in many cases prefer dead trees in specific stages of decay (Baker & Lacki 2006; Arnett & Hayes 2009; Fabianek *et al*. 2015). This knowledge has proven useful to biologists and land stewards working in forests managed for timber production or wildlife habitat, but there is not much known about the roosting ecology of many bat species. For example, a recent review of the effects of silviculture on bats contained no literature on species from South America (Law *et al*. 2016). Nevertheless, South American forests face extensive threats from several industries, climate change, wildlife diseases, and expansion of human settlements, and an understanding of the ecologies of bat species in this region is needed to promote their conservation (Armesto *et al*. 1998; Bustamante & Simonetti 2005; Wilson *et al*. 2005).

More than 200 species of bats are found in South and Central America (Altringham 2011), with the highest diversity residing in tropical rainforests (Simmons & Cirranello 2019; Wilson & Mittermeier 2019). Although only three species inhabit the sub-Antarctic forests located in austral Chile and Argentina (Koopman 1967), there are few studies on the ecology of species in the region, especially in comparison to species in the Northern Hemisphere. However, data on habitat use and behaviors of southern species are no less sorely needed, as habitat loss is likely to profoundly affect bats in the region. Indeed, the amount of native, undisturbed forest in Patagonia is declining due to several factors linked to human activities, such as timber harvesting (Lorenzo *et al*. 2018). Further, introduced species have changed the structure and composition of many southern forests. For example, beavers (*Castor canadensis*) have greatly disturbed riparian habitats due to the construction of their dams (Schiavini *et al*. 2019), and non-native pine plantations have replaced native forests with monocultures (Braun *et al*. 2017; Franzese *et al*. 2017).

In addition to being challenged by habitat loss, bats of the southernmost forests (i.e., those of Tierra del Fuego) are challenged by cool and variable weather conditions and short summers, potentially making them particularly vulnerable to loss of critical habitats. Cool temperatures are especially challenging for females during gestation and lactation, when energetic demands are already high (Kurta *et al*. 1989; McLean & Speakman 1999; Kerth *et al*. 2001). Compounding difficulties in meeting energy budgets while reproducing in cold environments, insect activity decreases during periods of cold (Scanlon & Petit 2008), and short summers limit the time available to prepare for hibernation (Jooste *et al*. 2013; Johnson *et al*. 2017). Finally, lack of true darkness during mid-summer may inhibit foraging or expose foraging bats to predators (Rydell 1992). Thus, bats living in the southernmost forests of the world face numerous challenges that must be met through physiological adjustments, behavior, and habitat selection.

A growing body of literature shows that reproductive females use torpor to conserve energy during periods of cold (Willis *et al*. 2006a; Johnson & Lacki 2014; Besler & Broders 2019), although this behavior delays fetal development and reduces milk synthesis (Racey & Swift 1981; Wilde *et al*. 1999). The need to use torpor, and experiencing the associated costs, is likely reduced through selection of warm roosts and social thermoregulation (Willis & Brigham 2007). Thus, roosts suitable for reproductive females are an essential aspect of summer habitat for bats at high latitudes. The characteristics of these roosts, and bats’ thermoregulatory behaviors within them, are therefore essential starting points for conservation and management efforts in these regions.

Our goal was to describe the roosting ecology of two species inhabiting the sub-Antarctic forest of southern Tierra del Fuego: *Myotis chiloensis* (Chilean myotis) and *Histiotus magellanicus* (southern big-eared brown bat). Because there are no descriptions of thermoregulatory strategies for either species, our primary objective was to determine how both species respond to cold spring temperatures in the region. We hypothesized that because both species would be challenged by the cold, they would rely heavily on torpor and use deeper torpor on colder days. We further hypothesized that because of its smaller size, *M. chiloensis* would use torpor for more time during the day than *H. magellanicus*. A secondary objective was to describe the roosts used by each species, as there are no descriptions of their daytime habitats in the sub-Antarctic forests. Because both species have been found to roost in tree crevices and under bark (Mann 1978; Altamirano *et al*. 2017), we hypothesized that *M. chiloensis* and *H. magellanicus* would be found roosting in trees with similar habitat characteristics. Our data provide insights into the ecologies of the southernmost bats that are an important foundation for conservation.

## MATERIALS AND METHODS

### Study Area

Our study took place in Karukinka Natural Reserve in southern Tierra del Fuego (54° S, 68° W, Figure 1). Karukinka is managed by the Wildlife Conservation Society (WCS) and consists of approximately 300,000 hectares of *Nothofagus pumilio* (lenga) and *N. betuloides* (coigüe de Magallanes) forest (44.2%), peat bogs (34.8%), mountains (5.5%), steppes (1.1%), and lakes and rivers (0.7%) (Wildlife Conservation Society 2018). Elevations range from sea level at the southwest side of the reserve to 1200 m in the Fuegian mountain range. The region has a sub-Antarctic climate, receiving 600–800 mm of precipitation annually, and is typically characterized by cold temperatures. The coldest months occur during austral winter, June– August, when mean daily temperatures fall below 0 °C. Austral summer occurs from December– February, when mean daily temperatures are typically < 10 °C.

**Figure 1.**
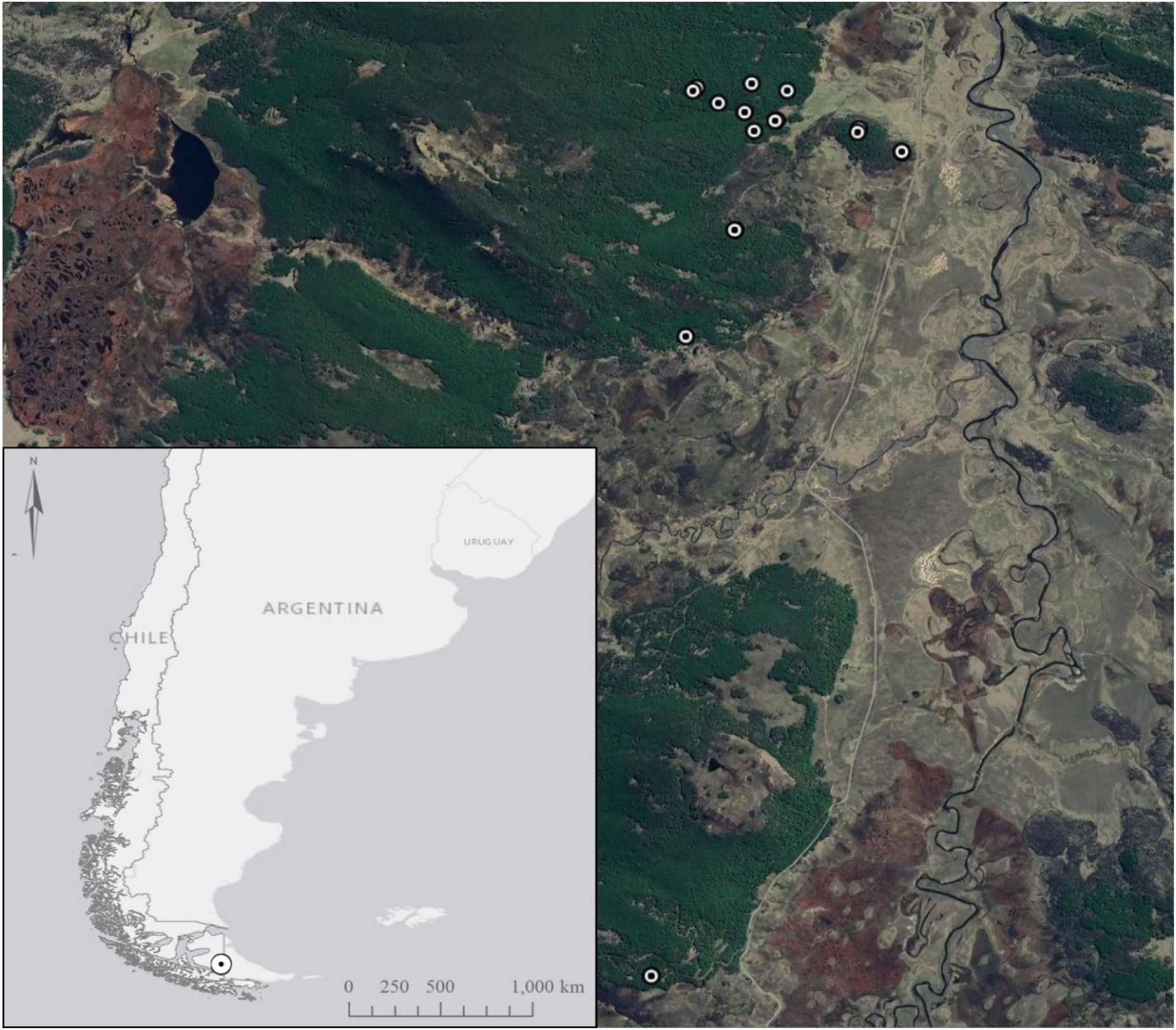
Satellite photograph of the study area within Karukinka Natural Reserve in Tierra del Fuego, Chile. Bullseyes show locations of day-roosts found by radio-telemetry within the study area. Because some roosts were located in close proximity to others, not all roosts are visible. Inset bottom left, a map of southern South America with the location of Karukinka Natural Reserve denoted with a bullseye.

During the 1990s, the Trillium timber company purchased the land that is now known as the Karukinka Natural Reserve for timber production, but the enterprise failed, and the land was donated to the WCS by Goldman Sachs in 2004. Today, Karukinka is a protected reserve managed by the WCS. Because Karukinka is home to some of the southernmost bat populations in the world, the reserve was recently recognized as an important area for the conservation of bats by the Latin American Network of Bat Conservation (Ossa *et al*. 2018).

### Study Species

We focused our research on *H. magellanicus* and *M. chiloensis*, two species believed to be non-migratory inhabitants of Tierra del Fuego (Koopman 1967). A third species, *Lasiurus varius* (cinnamon red bat) may also inhabit this region, based on an observation from the Magellan Strait. *Histiotus magellanicus* is the larger of the two species (forearm: 42.4–51.6 mm; weight: 12–22 g) and is found in native and non-native forests throughout central and southern Chile (Rodriguez-San Pedro *et al*. 2016; Díaz *et al*. 2019). This species roosts in holes and fissures within the largest decaying trees available in their stands (Altamirano *et al*. 2017), as well as underneath the loose bark of dead trees (Peña & Barria 1972). *Myotis chiloensis* (forearm: 33–42 mm; weight: 6–10 g) uses both anthropogenic and natural roosts and is often found in buildings (Ossa *et al*. 2010; Ossa & Rodriguez-San Pedro 2015). In tree roosts, this species is found in holes, fissures, and underneath the bark of dead trees (Mann 1978). *Myotis chiloensis* may have the most southerly distribution of any bat species, as it has been documented in sub-Antarctic *N. pumilio* forests in the Cordillera de Darwin (54°35’ S) (Ossa 2016).

### Capture and Data Collection

We captured and studied bats during the late spring, November and early December, of 2016 and 2017. Bats were captured using monofilament mist nets (Ecotone, Poland) set up inside *Nothofagus* forests. Mist nets were opened around sunset and remained open until midnight, when temperatures typically dropped below 0 °C, and when bat activity – monitored using a Pettersson D240X (Pettersson Elektronik, Sweden) – ceased. Captured individuals were measured and identified to species (Díaz *et al*. 2016), age, and reproductive condition. In order to find day-roosts, we affixed temperature-sensitive radio-transmitters (model LB-2XT, Holohil Systems, Canada) between the scapulae using surgical adhesive (Perma-Type, USA). Transmitters were individually calibrated by the manufacturer.

We searched for tagged bats each day using a receiver (Biotracker, Lotek, Canada) connected to a three-element yagi antenna. At each roost, we recorded habitat variables known to be important in other tree-dwelling species, and for which we had the equipment necessary to measure. Specifically, we recorded diameter at breast height using a diameter tape, roost tree and canopy height using a clinometer, and counted the number of live and dead trees > 10 and > 25 cm in diameter within 20 m of the roost tree. We determined the distance from each roost to the forest edge using Google Earth Pro (v. 7.3.2.5776 (64-bit)). Roost emergence counts were not performed due to a lack of personnel. Because we located relatively few roosts, we did not collect similar data from random trees for comparison and elected instead to compare the roosts between species. After locating forest stands where bats commonly roosted, we deployed datalogging receivers (RX800, Lotek, Canada) at high points overlooking those stands to detect and record skin temperatures (*T*_*sk*_) of tagged bats. Datalogging receivers were set to scan for each radio frequency for 30 seconds and to begin scanning for programmed frequencies at 5-min intervals. To document weather conditions during our study, we deployed a temperature datalogger (model MX2301A, Onset Computer Corp., Bourne, MA) inside a solar radiation shield within the forest, 350 m from the forest edge and at an elevation of 213 m. We programmed it to record air temperature (*T*_*a*_) at 15-min intervals throughout the duration of the study.

### Data Analysis

We performed all analyses in Program R v. 3.4.1 (R Core Team 2018). We compared diameter, tree height, distance to forest-non-forest edge, and the number of trees > 25 cm dbh within 20 m of roosts with Kuskal-Wallis tests. These analyses included both sexes of *H. magellanicus* and female *M. chiloensis* as distinct groups for comparison. We did not analyze the number of trees > 10 cm because while this measure represents the density of stems in the stand it provides less information on trees that may be large enough to be used by bats.

We reviewed *T*_*sk*_ data prior to analysis and removed days where data were not collected continuously between sunrise and sunset. We then summarized the minimum and variance in *T*_*sk*_ for each day of data and determined the amount of time (hrs) each bat spent in torpor. We used a mathematical equation to predict torpor onset (Willis 2007). This equation resulted in a torpor threshold value of 32 °C for both species. We considered all bats to be torpid when *T*_sk_ dropped below this threshold for two or more consecutive temperature readings. To test our hypotheses of the effect of species and daily weather on torpor use, we constructed three linear mixed-effects models (LME) using the lmer function in the lme4 package (Bates *et al*. 2015). Each model contained species and daily average *T*_*a*_ as fixed effects and the unique bat as a random effect but used different measures of *T*_*sk*_ (minimum and variance) or time torpid as the dependent variable. Sexes were pooled due to low sample sizes. In these models, minimum *T*_*sk*_ reflects the maximum depth of torpor bouts while variance in *T*_*sk*_ reflects how consistent *T*_*sk*_ was throughout the day. We calculated the *P* values for each fixed factor using the ANOVA function in the lmerTest package (Kuznetsova *et al*. 2017). We considered *P* ≤ 0.05 significant.

## RESULTS

We captured 30 adult *H. magellanicus* (14 male and 16 female) and 11 adult *M. chiloensis* (4 male and 7 female). Of these bats, we radio-tagged 17, including 11 *Histiotus* (6 male and 5 female) and six *Myotis* (1 male and 5 female). We were unable to locate four of these individuals, including two male *Histiotus* along with one male and one female *Myotis*. We tracked the remaining bats to a total of 17 day-roosts (Table 1). There were no differences in diameter (*K* = 0.26, *df* = 1, *P* = 0.61), height (*K* = 0.52, *df* = 1, *P* = 0.47), distance to forest edge (K = 3.21, *df* = 1, *P* = 0.47), or number of large trees within 20 m (K = 0.16, *df* = 1, *P* = 0.69) among *H. magellanicus* and *M. chiloensis* roosts. All roosts used by both species were in large *N. pumilio* trees (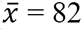; range = 46–108 cm), including three snags. Roosts varied widely in height (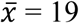; range = 7–27 m), ranging from trees that were snapped in half to those that extended > 10 m above the surrounding forest. Roosts were often found in dense stands (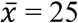; range = 6– 60 trees within 20 m) with few large trees (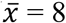; range = 0–32 trees > 25 cm within 20 m). We were unable to observe where bats roosted in trees as trees lacked visible cavities. Snags and even several live trees had obvious exfoliating bark where we presume bats roosted underneath. However, bats also roosted in live trees without obvious exfoliating bark, and we suspect that bats roosted in crevices within these trees. Roost trees were often located close to edges of forests and nearby peat bogs and steppes (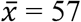; range = 0–675 m). Roosts located far from the edge of forest and non-forested habitats were often along forest gaps such as the edges of abandoned roads used for timber harvest (Figure 2A), stands where beavers had killed many of the trees in the overstory (Figure 2B), and areas of storm blowdown. Roosts were also consistently located at low elevations (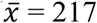; range = 162–275 m) near the base of large slopes.

**Table 1.**
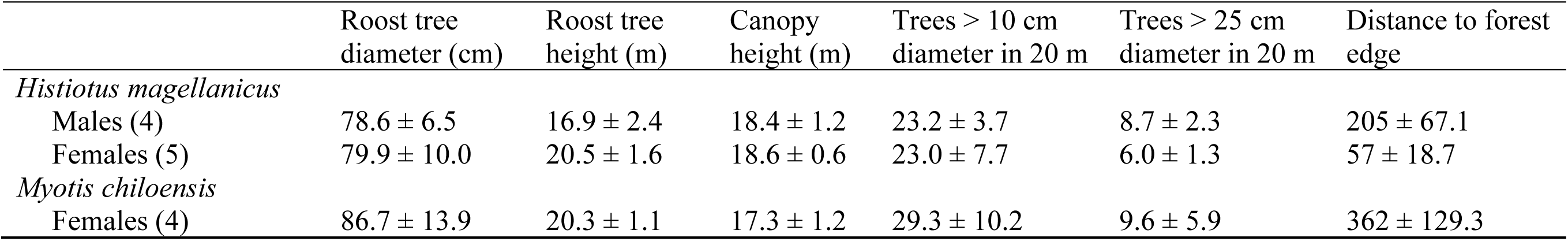
Characteristics of day-roosts used by *Histiotus magellanicus* and *Myotis chiloensis* (sample sizes in parentheses) determined by radio-telemetry in Karukinka Natural Reserve, Tierra del Fuego, Chile. Data reported are means ± S.E.

**Figure 2.**
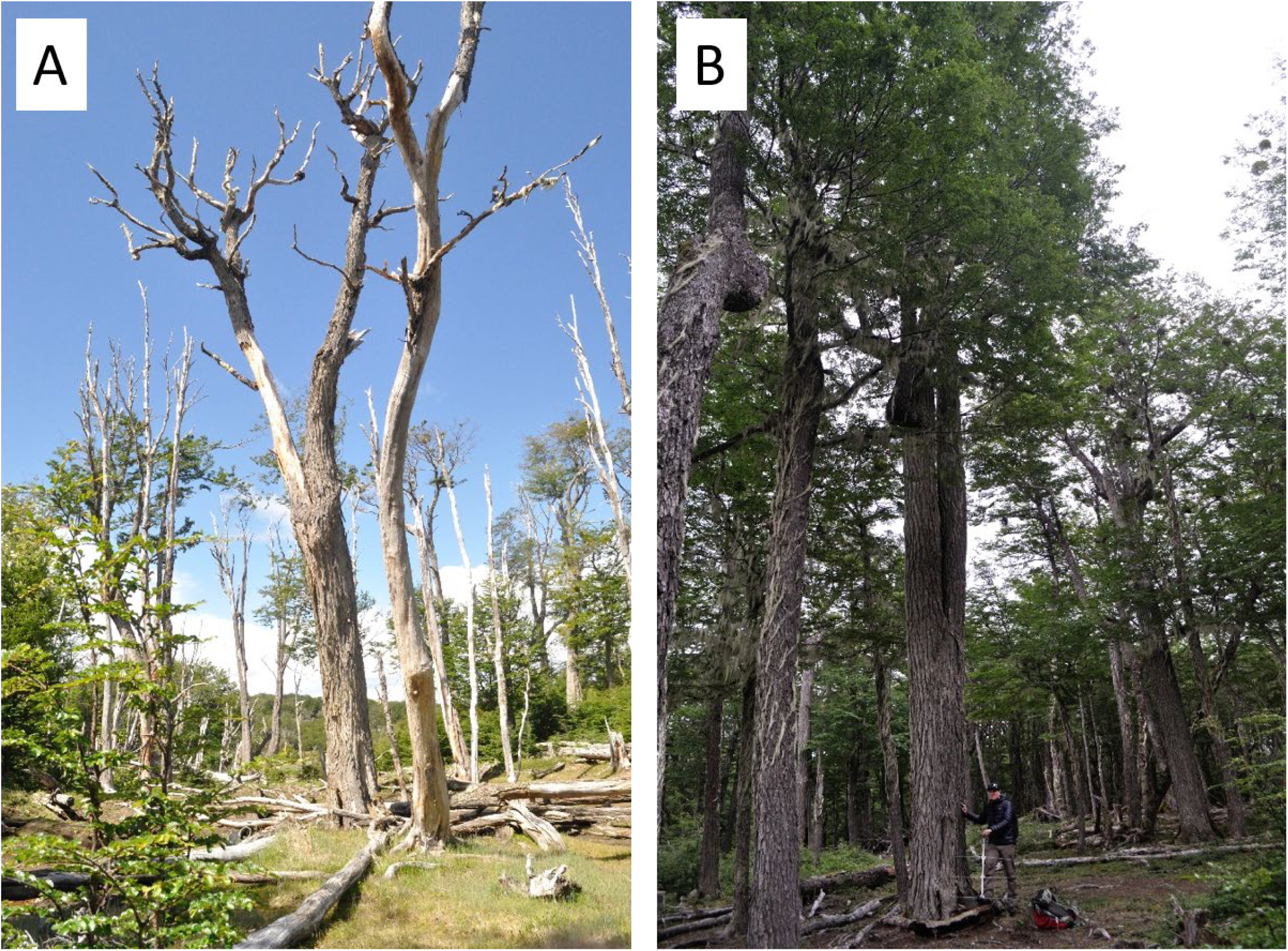
Photograph of *Nothofagus pumilio* used as day-roosts by *Histiotus magellanicus* (A) and *Myotis chiloensis* (B) in Karukinka Natural Reserve, Tierra del Fuego, Chile.

*Histiotus magellanicus* exhibited behaviors typical of bats with fission-fusion social groups. Of the 12 *Histiotus* roosts we located, three (25%) were used by more than one of our radio-tagged bats. These three roosts were separated by 214–547 m and were all located within the same forest stand located > 1 km from where all three bats were captured. These three bats were found roosting in the same tree on one occasion, and two were found roosting together on 80% of days we were able to locate at least one bat. We observed frequent roost switching among *Histiotus*, but only observed a single roost switch by *Myotis* (Table 2). None of the radio-tagged *Myotis* were tracked to a roost used by another tagged bat.

**Table 2:**
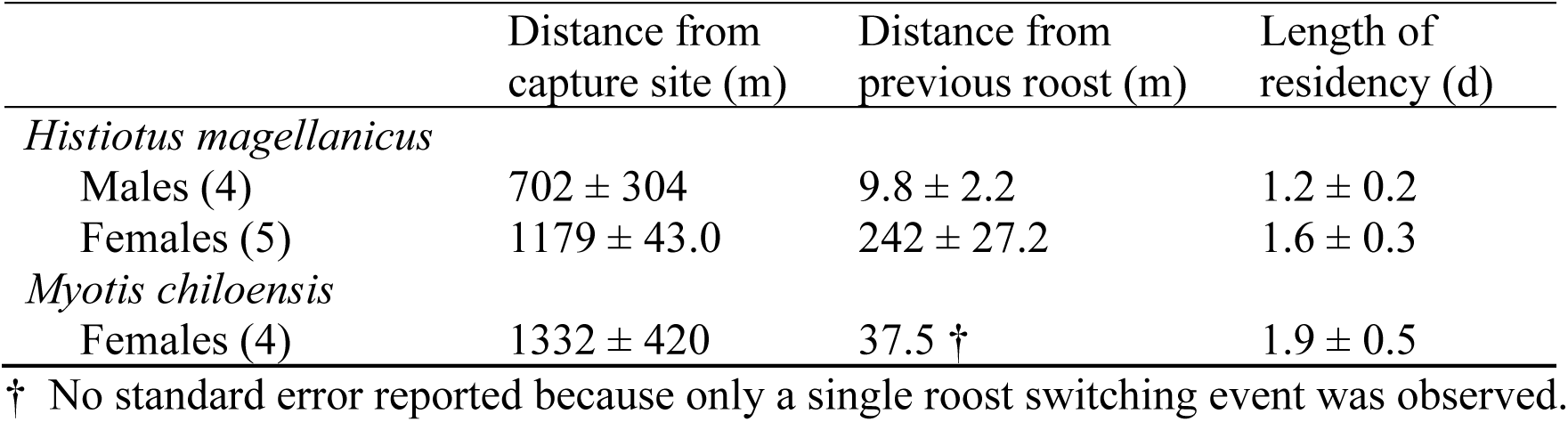
Distances traveled by radio-tagged *Histiotus magellanicus* and *Myotis chiloensis* between capture sites and day-roosts, distances travelled between day-roosts used during consecutive days, and number of days bats resided in the same roost before switching to a new location. Data reported are means ± S.E.

|  | Distance from<br>capture site (m) | Distance from<br>previous roost (m) | Length of<br>residency (d) |
| --- | --- | --- | --- |
| *Histiotus magellanicus* |  |  |  |
| Males (4) | 702 ± 304 | 9.8 ± 2.2 | 1.2 ± 0.2 |
| Females (5) | 1179 ± 43.0 | 242 ± 27.2 | 1.6 ± 0.3 |
| *Myotis chiloensis* |  |  |  |
| Females (4) | 1332 ± 420 | 37.5 † | 1.9 ± 0.5 |

† No standard error reported because only a single roost switching event was observed.

We successfully collected *T*_*sk*_ data during 81 full days (sunrise–sunset). This included 52 days of data collected from eight *H. magellanicus* (4 males and 4 females) and 29 days collected from four *M. chiloensis* (all female). Bats used torpor on 89% of days (*n* = 72). Bats spent an average of 11.7 hrs in torpor each day, with torpor bouts often lasting the entire day and with *T*_*sk*_ frequently dropping to a few degrees warmer than ambient temperatures on cold days (Figure 3). Bats used deeper torpor (i.e., had lower *T*_*sk*_) on colder days, as demonstrated by the significant positive effect of average *T*_*a*_ on minimum *T*_*sk*_ (*b* = 2.03, *t* = 4.77, *P* < 0.05) (Figure 4). Bats also spent more time in torpor on colder days (*b* = −1.52, *t* = −3.84, *P* < 0.05) but exhibited more variable *T*_*sk*_ on colder days as a result of skin temperatures following daily patterns in ambient temperature (*b* = −6.70, *t* = −3.63, *P* < 0.05). There was no significant difference in minimum *T*_*sk*_ (*b* = −3.88, *t* = −0.85, *P* = 0.41), time torpid (*b* = 1.62, *t* = 0.37, *P* = 0.72 or variance in *T*_*sk*_ (*b* = 20.71, *t* = 1.22, *P* = 0.25) between species (Figure 5).

**Figure 3.**
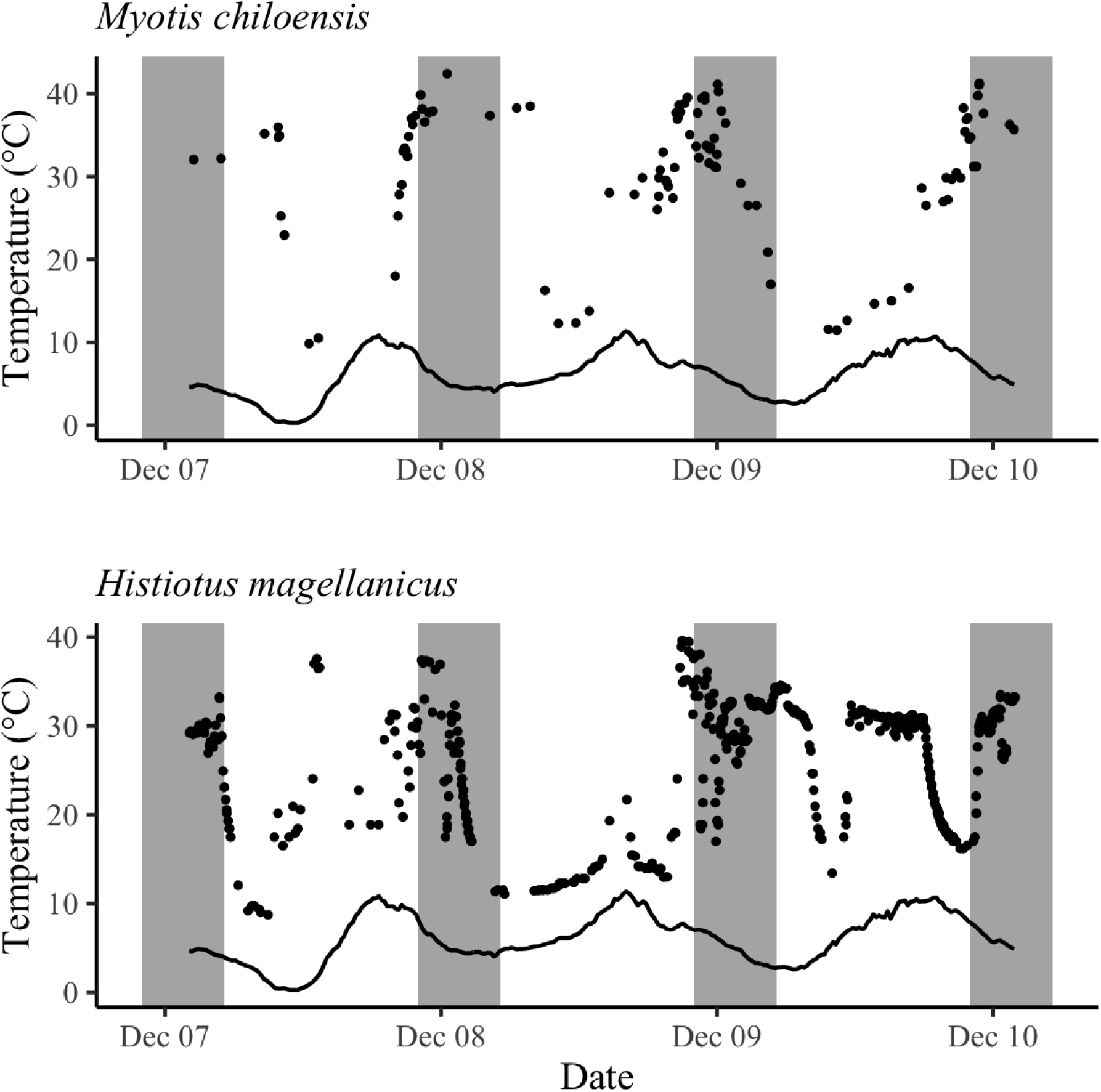
Skin temperatures (black circles) of *Histiotus magellanicus* and *Myotis chiloensis*, determined by radio-telemetry, and ambient air temperature (solid line) collected in December 2017 within our study area in Karukinka Natural Reserve, Tierra del Fuego, Chile. Hours between sunset and sunrise and contained within shaded areas of the graphs.

**Figure 4.**
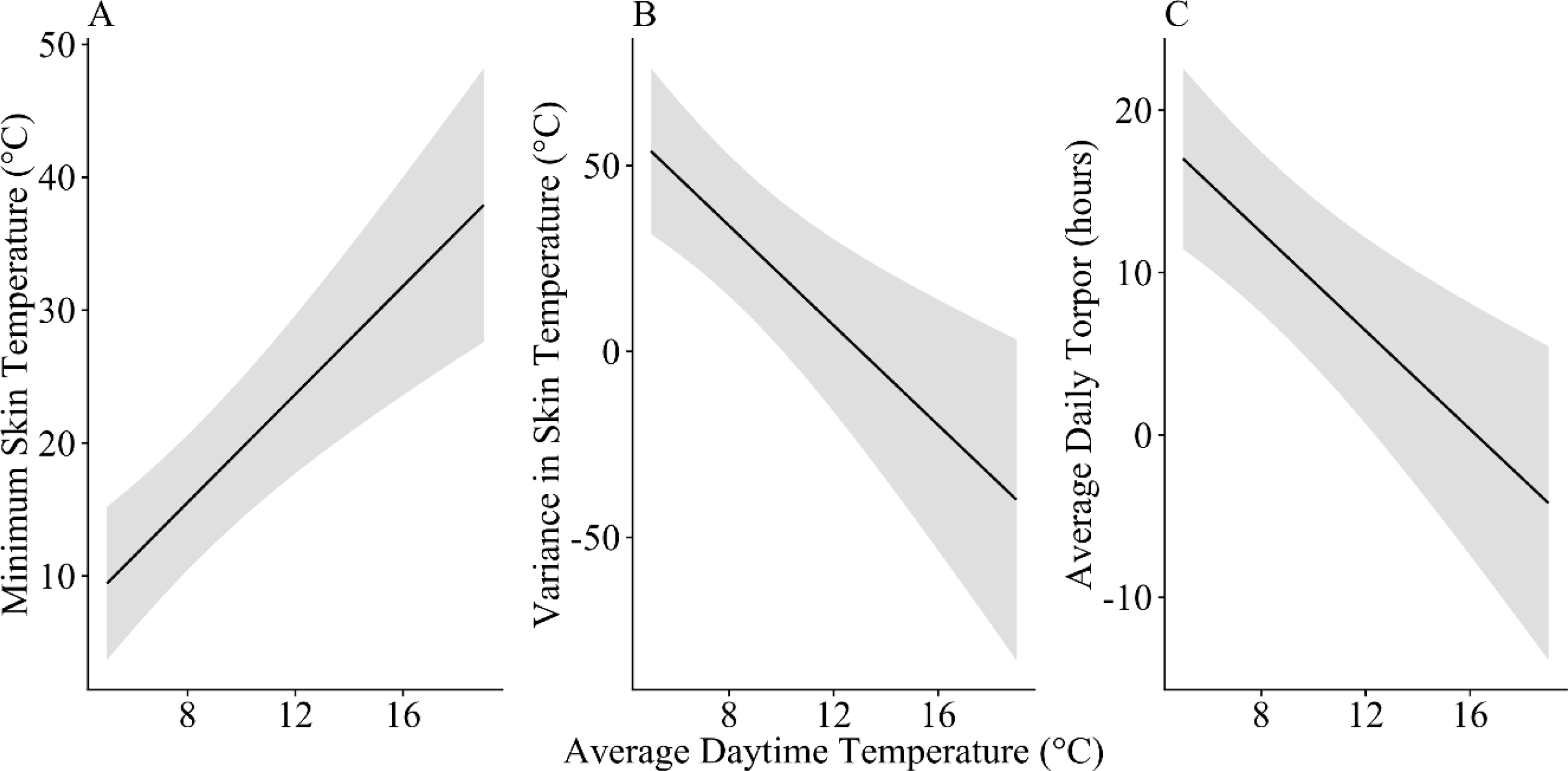
Predicted values of minimum skin temperature (A), variance in skin temperature (B), and time spent in torpor (C), and associated 95% confidence intervals (gray), of *Myotis chiloensis* and *Histiotus magellanicus* in relation to average daytime temperature.

**Figure 5.**
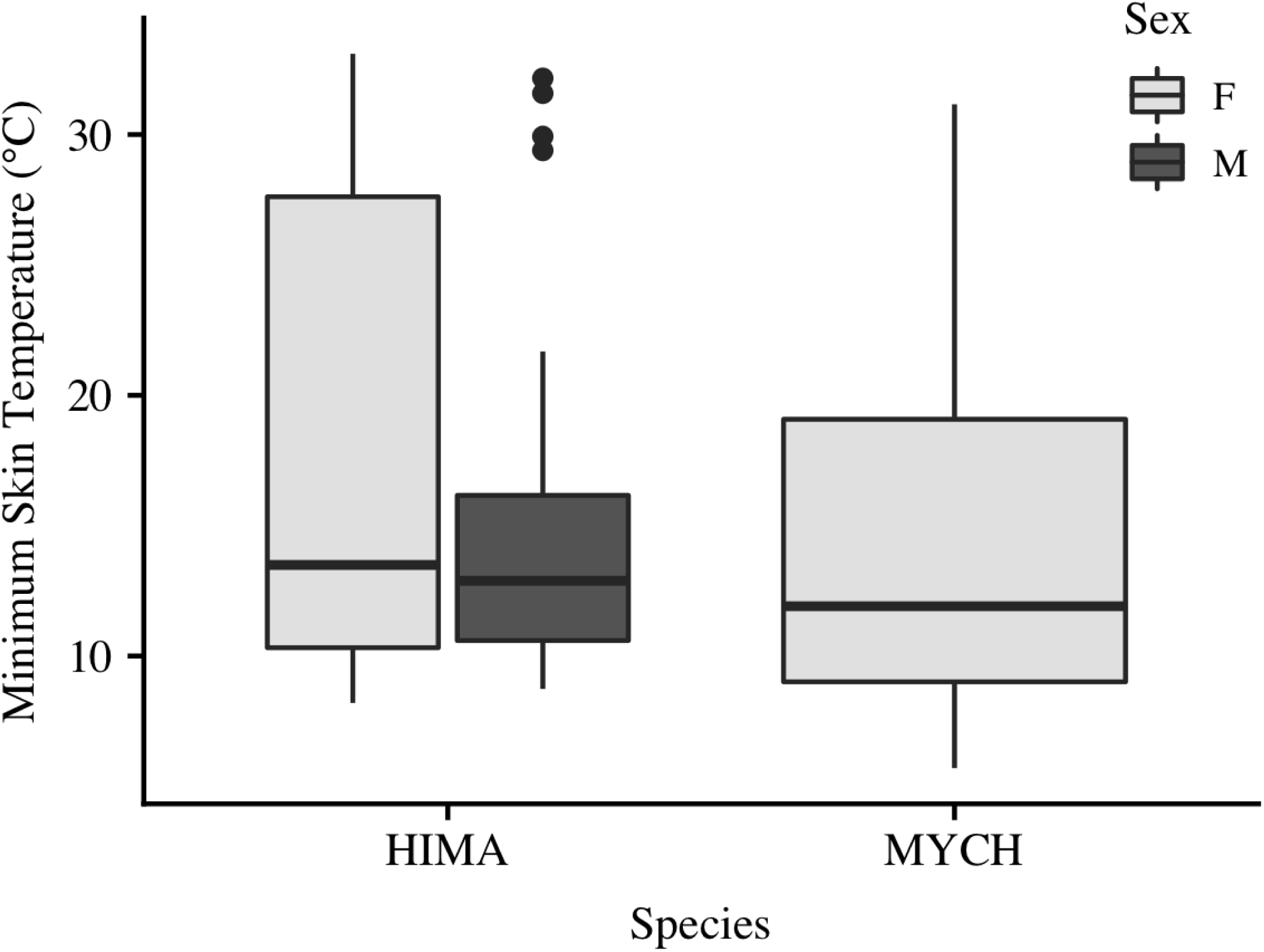
Minimum daily skin temperature did not differ between *Myotis chiloensis* (MYCH, *n* = 4) and *Histiotus magellanicus* (HIMA, *n* = 8) during austral spring (November and December) of 2016–2017 at the Karukinka Natural Reserve, Tierra del Fuego, Chile. Although sex was not a factor in our analysis, data for male and female *Histiotus* are graphed separately. In each box, medians are shown with a black line and observations outside 1.5 times the interquartile range are denoted with a closed circle.

## DISCUSSION

We provide the first descriptions of tree roosts and thermoregulation of *Myotis chiloensis* and *Histiotus magellanicus* inhabiting the southernmost forest of the world. Both species roosted in large *Nothofagus pumilio* trees at low elevations, frequently located along forest edges and gaps created for roads, wind storms, and beaver dams. As we hypothesized, the cold spring temperatures in the region caused both species to use torpor frequently, with both species relying on torpor more on colder days. However, contrary to our expectations, we found no difference in torpor use between *H. magellanicus* and the smaller *M. chiloensis*. These findings show the cold climate of southern Patagonian forests place small, insectivorous bats under energetic stress that bats meet through use of torpor and likely habitat selection as well.

The frequent use of torpor we documented in Tierra del Fuego is similar to that of male *M. daubentonii* (Daubenton’s bat) during early summer in the northern hemisphere, where low abundance of insect prey during this time makes it difficult for insectivorous bats to meet their energy budgets (Becker *et al*. 2013). Difficulty meeting energy budgets is likely to be more pronounced for females than males during spring, when females of species such as *M. chiloensis* and *H. magellanicus* become pregnant. In other species, pregnant females have been found to respond to this energetic challenge by using torpor to save energy and possibly delay parturition until favorable summer conditions arrive (Racey & Swift 1981; Willis *et al*. 2006a; McAllan & Geiser 2014). However, data from *M. lucifugus* (little brown myotis) suggest that delaying parturition past a certain point may influence juvenile survival, as bats born later in the year have lower first-year survival rates (Frick *et al*. 2010). Thus, torpor, while an important mechanism for saving energy during periods of cold, may come at a cost for some species, as it has been shown to slow fetal development, possibly reducing reproductive success (Dzal & Brigham 2013).

The climate of Tierra del Fuego appears to necessitate the use of frequent, and often deep (i.e., *T*_*sk*_ < 10 °C), torpor in both *M. chiloensis* and *H. magellanicus* despite the costs. Habitat selection may offset some of the need to use torpor, or, conversely, facilitate the use of torpor. In a study of insectivorous bats in Europe, Otto and colleagues (2016) found that in three similar sized bat species, species that used torpor more selected cooler roost than species that used torpor less. The similar skin temperatures and torpor patterns we observed for *M. chiloensis* and *H. magellanicus* in Tierra del Fuego suggests that roosts used by these species have similar microclimates. However, we did not collect data on roost microclimates or social group sizes, preventing direct comparison of habitats and microclimates between our study species. Furthermore, the lack of statistical difference we observed between species may be a result of small sample sizes or inclusion of male *H. magellanicus* in the analysis.

Given the costs of torpor use during reproduction, we hypothesize that both species select roosts that minimize thermoregulatory costs. We base this hypothesis on the observation that both species always roosted in large trees often located in forest gaps or along forest edges. Large diameter trees may offer larger cavities for bats, providing room for larger social groups, which can have a greater influence on microclimates within the roost than the characteristics of the roost tree (Willis *et al*. 2006b; Willis & Brigham 2007). Roost tree height can also affect roost microclimates, as demonstrated in a study of *Nyctalus noctula* (common noctule) roosts, which were warmer at higher roost heights due to increased solar exposure (Ruczynski 2006). High solar exposure and warmer roost microclimates can also be gained by selecting roosts with relatively open canopies (Bergeson *et al*. 2018). We did not measure solar exposure in our study, but it is likely that roosts located along edges and gaps benefited from more sunlight than trees in the forest interior (Figure 2).

Although our description of roosting habitat of *M. chiloensis* and *H. magellanicus* is based on a small sample that precludes a comparison of used to available trees, these data provide important insights into the ecology of these species in an area where few data are available. Indeed, relatively few studies exist that describe the roosts of these species. The most detailed description of *H. magellanicus* roosts come from Andean temperate forests far to the north (39° S latitude) (Altamirano *et al*. 2017). There, the species was found roosting within cavities in trees that were larger in diameter than surrounding trees, suggesting legacy structures are important to these bats. These results mirror our own, and while we did not measure available trees, all of our roosts were > 46 cm in diameter, and often > 100 cm. Similarly, although the *M. chiloensis* is known to roost in trees (Mann 1978; Ossa & Rodriguez-San Pedro 2015), descriptions of those trees are rare. We found *M. chiloensis* using similar roosts as *H. magellanicus*, and although our sample is limited, these results highlight the importance of large trees as habitat for both species.

The data presented in this study describe the thermoregulation and habitat use of the southernmost bats during late spring in southern Tierra del Fuego. Both *H. magellanicus* and *M. chiloensis* roosted exclusively in large trees where they used torpor nearly every day, including regular bouts dropping below 10 °C. Thus, these tree-dwelling species live in a thermally challenging environment where they roost in large trees that may buffer them from these conditions. We encourage further efforts to describe habitat use of bats in this region, and throughout the southern temperate and sub-Antarctic zone, so that important habitats may be identified and conserved.

## AKNOWLEDGEMENTS

This project was supported by The Rufford Foundation (Rufford Small Grant (10502-1 and 23042-2) and a grant from the Ohio University Research Committee. We thank Servicio Agricola y Ganadero (SAG) for capture permits in Tierra del Fuego (Res Ex: 1253/2016 and 4924/2017), Juan Carlos Aravena from the Instituto de la Patagonia for his help, and the Wildlife Conservation Society in Chile for allowing us to conduct research at Karukinka Natural Reserve and for their help with field work.

